# Anthropogenic habitats shape gut microbiome composition in Southern Indian bats

**DOI:** 10.64898/2025.12.23.695677

**Authors:** Vikram Iyer, B.R. Ansil, Darshan Sreenivas, Avirup Sanyal, Uma Ramakrishnan, Balaji Chattopadhyay

## Abstract

**Background:** Anthropogenic habitat modification and associated resources can exert selection pressures on wildlife and their microbiomes, altering their diversity, resulting in homogenization and making them resemble the human microbiome. Diet is an important predictor of the microbiomes of volant vertebrates, mainly for birds, but for bats, results remain inconclusive. In spite of India’s exceptional bat diversity, there is little understanding of how their microbiomes respond to anthropogenic habitats. Therefore, we investigated the trends of taxonomic and functional diversity and their relationships with host feeding-guild and phylogeny for six wide-ranging bat species across six anthropogenically modified sites in Southern India by generating 16S barcode sequences from their fecal samples.

**Results:** Eubacteria dominated samples with diet-specific taxonomic composition. Frugivore microbiomes contained large proportions of Cyanobacteria, possibly sourced from consumed plant matter or polluted drinking-water sources, and Lactobacillales dominated insectivore microbiomes, while Gammaproteobacteria were abundant regardless of host feeding guild. We found human pathogens in our samples possibly transferred from polluted water to the guts of bats foraging in nearby areas. We observed diet-specific taxonomic, phylogenetic, and functional composition. However, functional composition incorporating abundances displayed a high degree of overlap across feeding-guilds, suggesting functional homogenisation of microbiomes across feeding guilds, possibly due to anthropogenicity. Sample-wise diversity indices were significantly different with respect to diet only when samples from the same roost were not pooled together. However, in all cases, microbiomes from the same diet types displayed significant taxonomic and phylogenetic similarity to each other. Lastly, we observed limited concordance of microbiome diversity with chiropteran phylogeny.

**Conclusions:** Because of these potential signatures of pollution on bat microbiomes such as pathogens, we recommend their monitoring, especially because Cyanobacteria play a known role in bat and human disease. Our study is one of the first to study microbiome composition and function from bat species common around human inhabitation in South India, and establishes baselines in this region.

## Background

The animal host and its microbiome interact and shape each other. While the animal microbiome is shaped by the phylogenetic constraints of its host and host-dependent factors like diet, the relative importance of these two factors is clade-dependent across a wide range of animal taxa [1,2]. In corals, microbiome richness and composition vary across tissues with different levels of phylosymbiosis [3]. In mammals, phylosymbiosis primarily determines microbiome composition [4], although diet plays a role as well [1]. Specifically in apes [5] and hominids [6], gut microbiome similarity closely follows phylogenetic similarity. Such phylosymbiosis is stronger in internal compartments like the gut microbiome [7], and strong phylosymbiosis is observed across the gut microbiomes of vertebrates except in bats and birds which converge with each other, possibly due to adaptations to flight [8].

Host traits influence the gut microbiome as well, both in terms of the host’s feeding guild [9,10] as well as the specific food items consumed [11–14]. In the absence of phylosymbiosis as a dominant factor, a strong association between gut microbiome composition and diet is found in birds [9,15]. Results from bats (order Chiroptera), on the other hand, are [16,17] inconclusive. In some cases, while phylosymbiosis can be the dominant factor shaping bat microbiomes [18], in others, diet can outweigh phylogeny [19]. Functionally, bat microbiomes are convergent also based on host diet [20,21], although this may or may not be accompanied by taxonomic convergence [21,22].

Differential diet composition between anthropogenically modified habitats (like urban areas and intensively cultivated areas) and pristine habitats can exert large selection pressures on wildlife resulting in altered community composition [23–25]. Moreover, the food available to hosts can shape their microbiomes [26–29]. Recent studies suggest that gut microbiomes of urban mammals may be functionally diverse [30] and taxonomically homogeneous [29,30]. The effects of these pressures on urban microbiomes are poorly studied for non-charismatic taxa such as bats, which are exceptionally diverse, both taxonomically as well as functionally.

India has a rich diversity of bats, comprising 133 recognised species [31] across a range of dietary habits, feeding strategies (the ability to echolocate), and body sizes. Several wide-ranging Indian bat taxa are urban residents such as *Pteropus medius* [32–34], *Cynopterus sphinx* [35], and *Hipposideros speoris* [36] and are likely to display signatures of their anthropogenically modified habitat. This is especially likely because captivity and anthropogenic pressure both change animal microbiomes in predictable ways, often making them resemble human microbiomes [37–40]. Although anthropogenic modifications are known to affect Indian bats species’ ecology and behaviour [31,41,42], how they affect their microbiomes is poorly understood.

We investigated the gut microbiomes of a poorly understood urban bat community in Southern India. We collected the guano of six different bat species belonging to two different feeding guilds from several roosts across sites in and around Bengaluru in Southern Karnataka (Figure 1, Table 1), generated bacterial 16S barcode sequences and addressed the following questions:

1. What is the taxonomic composition of the guano microbiomes of bats?
2. How does this microbiome diversity vary compositionally and functionally across bat host taxa and diet?
3. Does microbiome composition correlate with host phylogeny?

**Figure 1:**
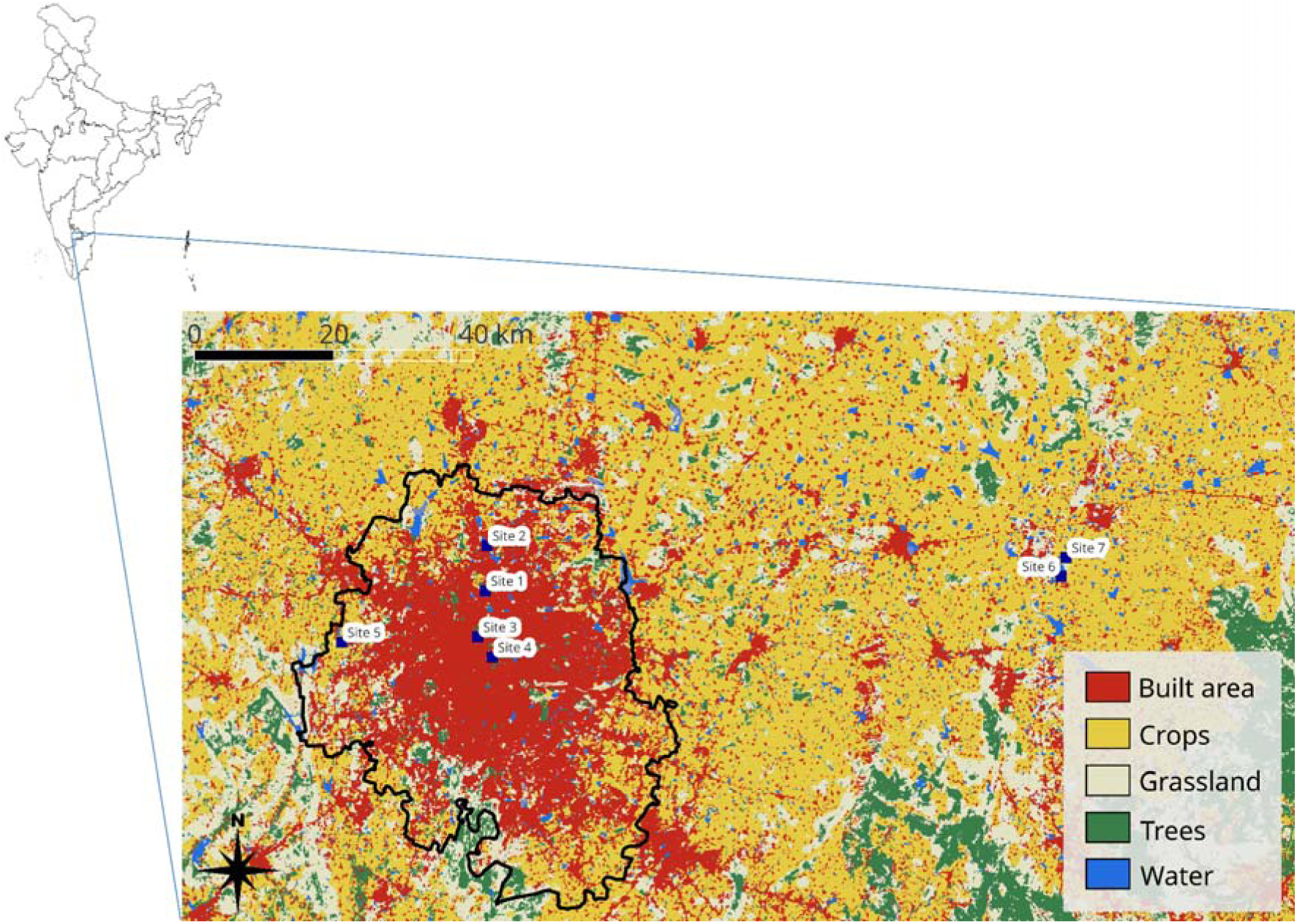
Map of the study area and sampling sites. The expanded inset map displays our study sites along with land-use types. The black outline represents the limits of Bengaluru district in South India.

**Table 1:** Details of all the unpooled samples analysed in this study.

| Sample ID | Species | Location | Collection date | Diet |
| --- | --- | --- | --- | --- |
| A1a | *Cynopterus sphinx* | Site 1 | 13.09.2021 | frugivore |
| A1b | *Cynopterus sphinx* | Site 1 | 22.10.2021 | frugivore |
| A1c | *Cynopterus sphinx* | Site 1 | 16.11.2021 | frugivore |
| A1d | *Cynopterus sphinx* | Site 1 | 23.11.2021 | frugivore |
| A1e | *Cynopterus sphinx* | Site 1 | 02.12.2021 | frugivore |
| A1f | *Cynopterus sphinx* | Site 1 | 09.12.2021 | frugivore |
| A1g | *Cynopterus sphinx* | Site 1 | 15.12.2021 | frugivore |
| A1h | *Cynopterus sphinx* | Site 1 | 04.01.2022 | frugivore |
| A1i | *Cynopterus sphinx* | Site 1 | 17.01.2022 | frugivore |
| A1j | *Cynopterus sphinx* | Site 1 | 18.01.2022 | frugivore |
| B1a | *Pteropus medius* | Site 2 | 16.09.2021 | frugivore |
| B2a | *Pteropus medius* | Site 3 | 23.03.2022 | frugivore |
| B2b | *Pteropus medius* | Site 3 | 25.05.2022 | frugivore |
| B2c | *Pteropus medius* | Site 3 | 02.05.2022 | frugivore |
| B2d | *Pteropus medius* | Site 3 | 11.05.2022 | frugivore |
| B3a | *Pteropus medius* | Site 4 | 26.03.2022 | frugivore |
| C1a | *Hipposideros speoris* | Site 5 | 26.11.2022 | insectivore |
| C1b | *Hipposideros speoris* | Site 5 | 27.01.2023 | insectivore |
| C2a | *Hipposideros speoris* | Site 6 | 05.02.2023 | insectivore |
| C2b | *Hipposideros speoris* | Site 6 | 08.06.2023 | insectivore |
| C2c | *Hipposideros speoris* | Site 6 | 09.06.2023 | insectivore |
| D1a | *Lyroderma lyra* | Site 6 | 08.06.2023 | insectivore |
| D1b | *Lyroderma lyra* | Site 6 | 09.06.2023 | insectivore |
| D2b | *Lyroderma lyra* | Site 7 | 05.02.2023 | insectivore |
| E1a | *Pipistrellus* | Site 1 | 22.11.2022 | insectivore |
| E1b | *Pipistrellus* | Site 1 | 23.11.2022 | insectivore |
| E1c | *Pipistrellus* | Site 1 | 12.01.2023 | insectivore |
| E1d | *Pipistrellus* | Site 1 | 19.05.2023 | insectivore |
| E1e | *Pipistrellus* | Site 1 | 20.05.2023 | insectivore |
| E1f | *Pipistrellus* | Site 1 | 24.05.2023 | insectivore |
| E1g | *Pipistrellus* | Site 1 | 30.05.2023 | insectivore |
| E1h | *Pipistrellus* | Site 1 | 20.06.2023 | insectivore |
| E1i | *Pipistrellus* | Site 1 | 26.05.2023 | insectivore |
| E1j | *Pipistrellus* | Site 1 | 27.05.2023 | insectivore |
| E1k | *Pipistrellus* | Site 1 | 28.05.2023 | insectivore |
| E1l | *Pipistrellus* | Site 1 | 29.05.2023 | insectivore |
| E1m | *Pipistrellus* | Site 1 | 31.05.2023 | insectivore |
| F1a | *Hipposideros lankadiva* | Site 6 | 08.06.2023 | insectivore |
| F1b | *Hipposideros lankadiva* | Site 6 | 14.07.2023 | insectivore |

## Results

### Structural and functional composition and diversity

We observed an overwhelming dominance of Eubacteria in the faecal microbiome of all the bat roosts that were sampled (Figure 2a). Most samples returned 100% eubacterial sequences except for *Pipistrellus,* which returned up to 1.55% archeal sequences (Supplementary figure S2a). At finer taxonomic scales, diet-specific microbiomes were observed. Frugivore microbiomes contained large proportions of Cyanobacteria, while insectivore microbiomes contained large proportions of Lactobacillales (Supplementary figure S2b – c) belonging to the Firmicutes D group. Gammaproteobacteria were abundant in most samples regardless of diet type. Additionally, some samples contained substantial abundances of Actinomycetales and Bacteroidales, also regardless of diet type (Figure 2a).

**Figure 2:**
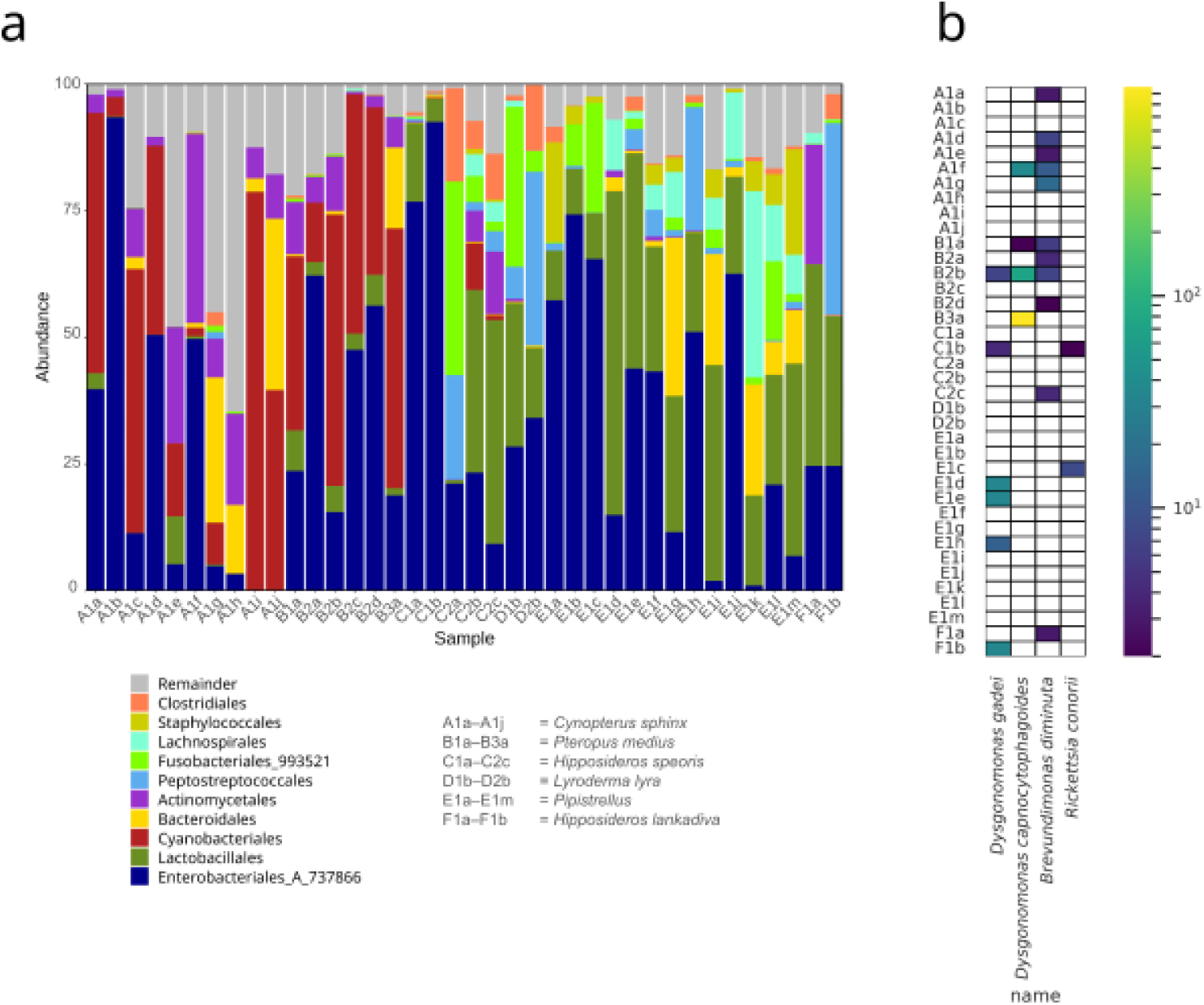
The taxonomic composition of bat microbiomes. a: Stacked bar plots displaying the proportion of various taxa in each sample at the order level. The 10 most abundant taxa at each level are displayed by name. The x-axis represents sample codes and the y-axis displays percentages. b: A heatmap of the pathogens detected in the samples. Colours indicate the number of ASVs obtained from each sample which correspond to a pathogen included in the MBPD pathogen database. Zero reported ASVs are shown in white.

We found four species of human pathogens in our samples which are known to occur in bats (Figure 2b), namely *Brevundimonas diminuta, Dysgonomonas capnocytophagoides, Dysgonomonas gadei,* and *Rickettsia conorii*. Most such pathogens were observed in the fecal microbiome of frugivores (10 out of 16 frugivore samples reported at least one pathogen), compared to insectivores (8 out of 23 samples). *Dysgonomonas capnocytophagoides,* the pathogen with the maximum ASVs (967 ASVs) represented, was found from a single sample of the frugivorous *Pteropus medius.* All other pathogens were represented with < 100 ASVs in each sample.

PCoA plots confirmed diet specific similarity in microbiomes (Figure 3a, supplementary figures S3 – S4). In all cases, the first principal component separated out samples based on diet regardless of the distance metric used and whether samples were pooled based on roost. Taxonomic presences from both unweighted (Jaccard metric) as well as weighted by abundances (Bray – Curtis metric; Supplementary figure S3) methods revealed two distinct *Pipistrellus* clusters from the same roost. However, this pattern was not present when a distance metric incorporating phylogenetic distances (Figure 3a) was used. Microbiomes of samples from *Pipistrellus* species segregated into two taxonomically distinct clusters, but these clusters were not phylogenetically distinct. Functional analyses showed a similar pattern—PCoA plots using predicted enzyme (EC) presences in samples revealed that the first principal component partitioned samples based on diet regardless of whether pooling was done (Figure 3b, Supplementary figure S5).

**Figure 3:**
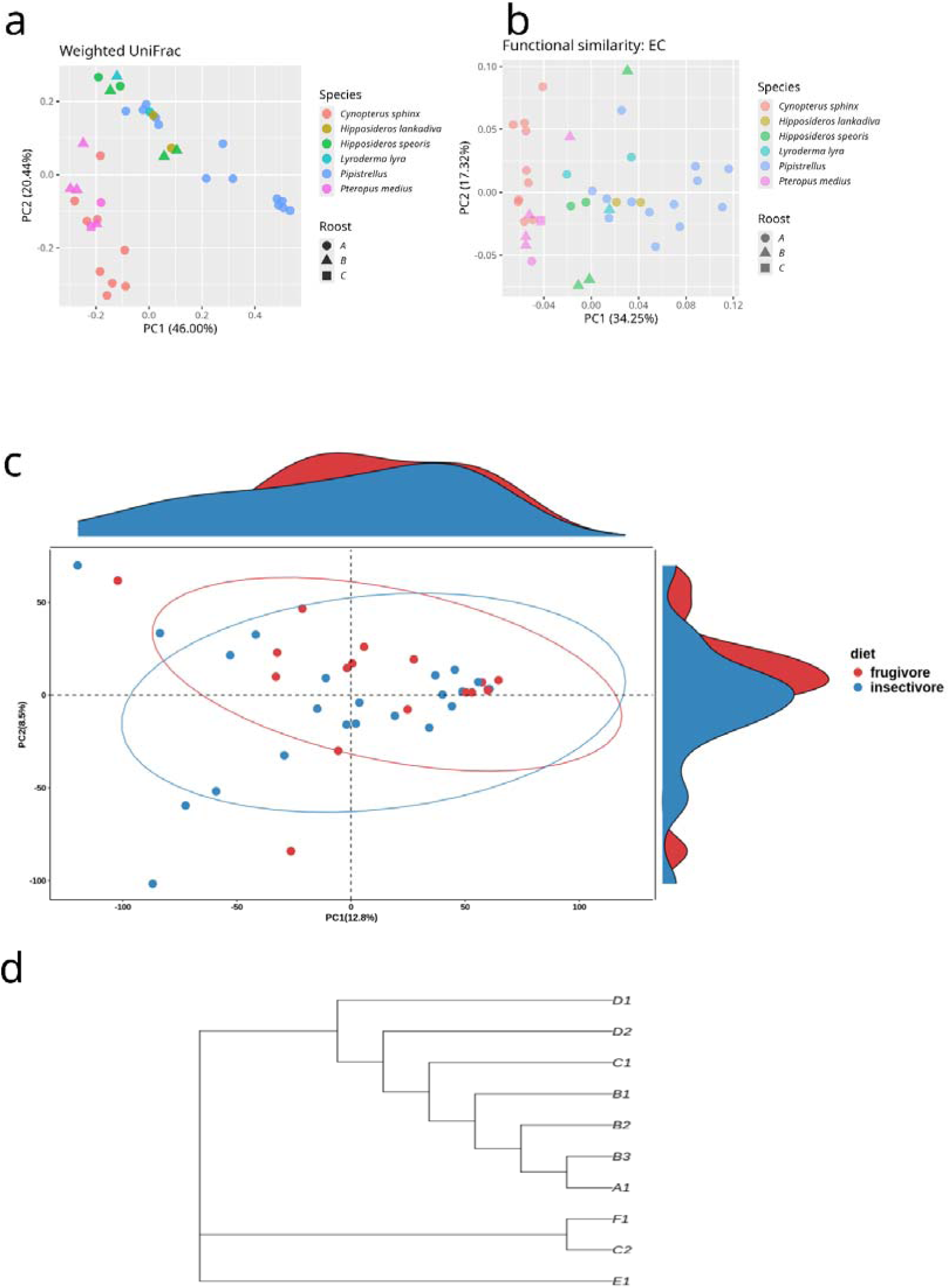
Compositional relatedness among samples. a: A Principal Coordinates Analysis plot of unpooled samples using the Weighted UniFrac distance metric and microbiome taxonomy and coloured by species identity. Each point represents a sample collected on a given date from a given roost. Points with the same colour and shape represent samples collected from the same roost on different dates. b: Principal Coordinates Analysis plot of unpooled samples using Jaccard distance metric and predicted enzyme abundances as per the Enzyme Commission (EC) and coloured by species identity. Each point represents a sample collected on a given date from a given roost. Points with the same colour and shape represent samples collected from the same roost on different dates. c: Principal Components Analysis plot of unpooled samples of predicted pathway abundances using KO pathway numbers. Ellipses indicate 95% confidence ellipses. Density plots are plotted on the axes. d: A neighbour-joining tree using pooled samples using the Weighted UniFrac distance metric.

Incorporating functional abundance information showed that bats with different diet habits have a high degree of functional overlap (Figure 3c, Supplementary figure S5). This was evident from the fact that the first two principal components explained 21.3% of the total variation in the data without pooling, and 44.4% of the total variation in the data after pooling. 95% confidence ellipses showed large amounts of overlap as well.

All three of the sample-wise diversity metrics used were significantly different between diet types before pooling (Figure 4) but were not significant after pooling (Supplementary figure S6). However, samples displayed significant taxonomic and phylogenetic relatedness to other samples of the same diet type as compared to samples of a different diet type regardless of the distance metric used to quantify this, and whether or not pooling was done (Figure 4, Supplementary figure S7 – S8). The fact that all frugivore samples belonging to the family Pteropodidae forming a clade as well corroborates this pattern (Figure 3d). Moreover, the *Pipistrellus* species was the only Yangochiropteran to segregate separately from its counterparts. Beyond this, however, we did not observe concordance of microbiome diversity with chiropteran phylogeny. There remained a polytomy at the root of the tree and Hipposideridae bat samples did not form a clade.

**Figure 4:**
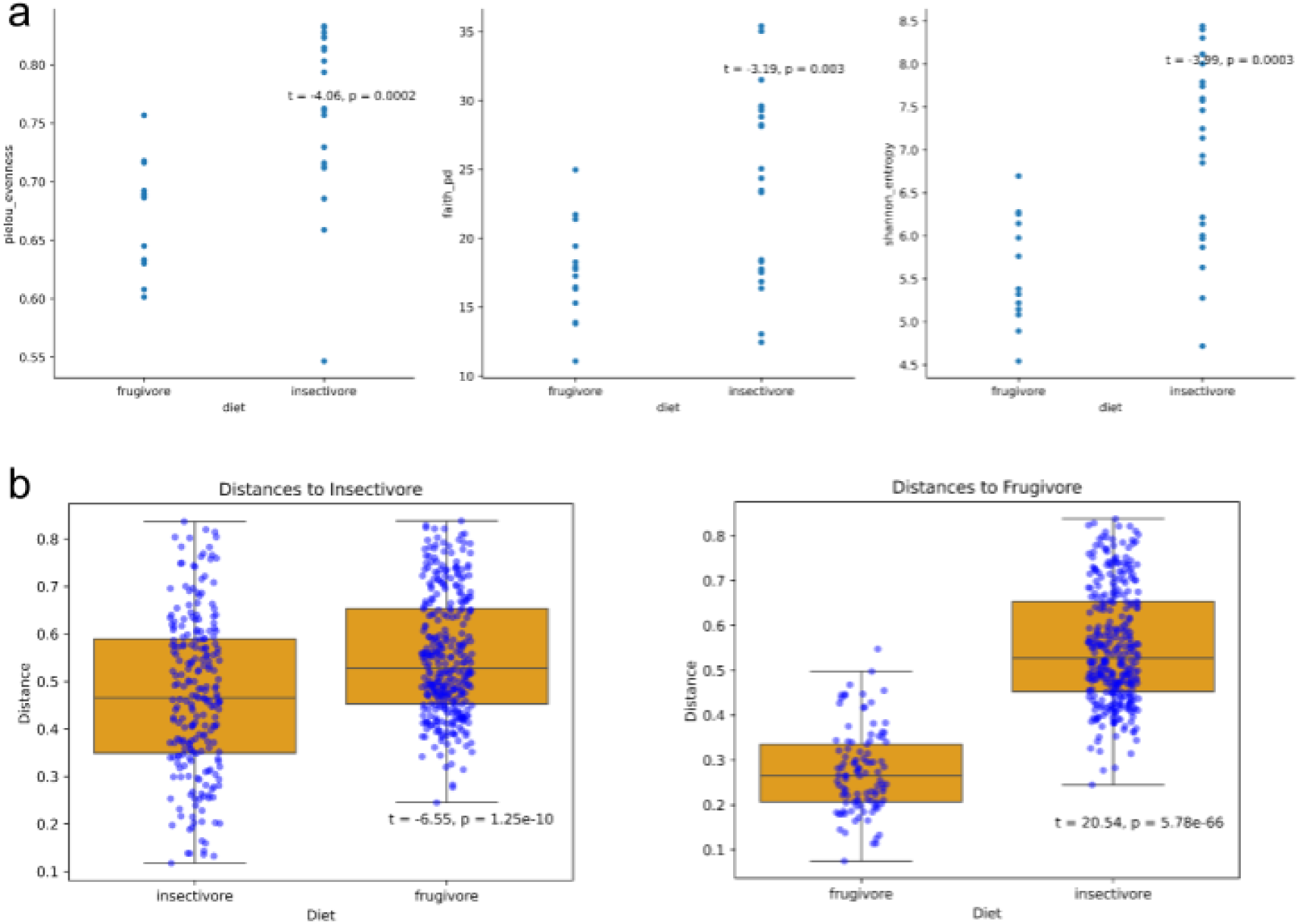
Alpha and beta diversity metrics for unpooled samples. Top: Strip plots of Pielou Evenness, Faith’s PD, and Shannon Entropy for each diet type. t and p values represent values from two-sided t-tests. Bottom: A boxplot of the distances of each datapoint to other data points of the same or different diet type. The Weighted UniFrac distance metric was used. All data points are overlaid in blue.

## Discussion

Urban habitats uniquely shape host habits that may leave characteristic footprints in their gut microbiome and reflect their ability to adapt. This study adds to the growing literature on bat microbiota and their determinants. We analysed the microbiomes of six tropical bat species from anthropogenically modified habitats across diet-types and observed homogenization of the microbiome reflecting the influence of human modification in shaping bat microbiome.

### Bat microbiomes are diet-specific

Our results confirm known knowledge about bats’ microbiomes while revealing novel diet-specific insights. Proteobacteria and Firmicutes are known to dominate in both the Old and New World bat microbiomes across diet-types [18,43] such as in nectarivorous [16,44,45], insectivorous [46,47], and frugivorous [16,46] species. In agreement, most of our samples reported large proportions of Gammaproteobacteria in a diet non-specific manner (Figure 2a). However, Firmicutes form a large proportion of the microbiomes of insectivores but not frugivores in our samples. Cyanobacteria have been previously reported in small proportions from certain life-history stages of a New World nectarivore bat species [44], and in large amounts from flying foxes in Guam [48,49]. One previous study from India that analysed guano samples from *Rousettus leschenaultii* also detected Cyanobacteria, albeit at lower abundances [50]. Our results show large proportions of Cyanobacteria from both frugivore species which are absent in all but two insectivore samples. These are likely to be not associated with life-history stage as our samples are roost-level averaged faecal samples collected over several months. Cyanobacterial toxins are known to be toxic to bats and their predators and have been associated with human illness as well [48]. Considering their notable presence in frugivorous bats from our study panel, we recommend detailed monitoring of their presence in Indian frugivore bats.

### Bat microbiomes are structurally and functionally similar

Microbes have high levels of functional redundancy, and independent taxonomic and functional segregation of microbial communities is possible [51]. Our results show that frugivorous and insectivorous bats are compositionally distinct in terms of microbiota taxonomy as well as phylogeny as evidenced by PCoA clustering (Figure 3a–b; Supplementary Figure S3 – S5) and distances within and across diet-types (Figure 4b, Supplementary Figure S7 – S8). However, quantitative diversity metrics between the two diet types are significantly different before pooling but not after. Some studies on bats report non-significantly different diversity metrics between diet types [21,45,50,52], but others report significant differences [16,17]. Our non-significant results after pooling could be an artefact of too few data-points. Studies looking at microbiome differences with varying life-history stages report non-significant differences in diversity metrics as well [52,53]. This could imply that while microbiomes can vary taxonomically and functionally across host taxa and the host’s lifespan, the limit of how much diversity an individual can support stays relatively constant.

Although we find that diet affects the functional types present in the microbiome (Figure 3b, Supplementary figures S5), integrating abundance information shows a high amount of functional overlap (Figure 3c, Supplementary figures S5). While only taxonomic [29] or taxonomic but not functional homogenisation of microbiomes [30] in urban regions is reported in other taxa, our results provide evidence of functional but not taxonomic homogenisation.

Sample-wise diversity metrics are relatively standardised across studies and there exist several tools for predicting functional abundances using 16S barcode sequences [54–56], but there is little standardisation on how functional annotation of barcode sequences is carried out. While these different tools fare comparably, their accuracy is low for non-human datasets [57] and we advocate caution in generalising from studies using different methods, especially for such non-human microbiome datasets.

### Anthropogenically modified habitats likely shape bat microbiomes

We highlight two possible sources for the Cyanobacteria present in frugivore fecal microbiomes. They might be sourced from consumed plant matter such as fruits or leaves. The near absence of this bacterial group in insectivorous bat microbiome (Figure 2a) supports this hypothesis. However, Cyanobacteria have not been consistently reported from other studies on microbiomes of frugivorous bats (Carrillo-Araujo et al., 2015; Dai et al., 2024), suggesting that diet alone may not account for their presence in fruit bat samples. An alternative and more plausible explanation is environmental exposure linked to anthropogenic pollution. Cyanobacteria from polluted water sources can be transferred to bats foraging or drinking in contaminated areas leading to dysbiosis [58]. Given that Cyanobacterial blooms are common in eutrophic water bodies [59], their dominance in the frugivore *Pteropus medius* and *Cynopterus sphinx* samples might be indicative of them using large, polluted water bodies (common in the study area) as a drinking-water source, thus reflecting additional exposure to such environmental pollution rather than dietary intake alone.

All four of the bacterial pathogens that we observed to be present within our study panel are human pathogens based on the MBPD database, and no plant or animal bacterial pathogens were obtained from our samples (Figure 2b). *Rickettsia conorii* is a widespread pathogen causing dog-tick spread Rickettsioses [60] and *Brevundimonas* sp. are being recognised as emerging opportunistic pathogens which have been so far neglected [61]*. Dysgonomonas* sp. can cause illness like bacteremia and are reported from a wide variety of tissues but are difficult to isolate clinically [62]. Strain-level taxonomic classification is often required for pathogenicity, and our samples lack this resolution. While this points towards the humanisation of the microbiomes of our samples, our results are preliminary in nature, and conclusively testing this requires extensive survey to analyse human microbiomes from our study region. Because our data are sequenced to an adequate depth (Supplementary figure S1) and only a few pathogen reads were reported in samples (Figure 2b), these are likely present in trace amounts. All our samples were collected from highly anthropogenically modified habitats, and observing human-infecting pathogens in our datasets points towards the possibility of the transfer of human microbiome to other mammals like bats in densely populated areas. Several of our samples were collected in Bengaluru, whose water quality is extremely poor across metrics [63–65] and the human-infecting pathogens in bats’ faeces might be sourced from these polluted water sources. We advise monitoring human pathogens in wildlife, especially because they may lead to microbiome dysbiosis and serious illnesses in wildlife. Additionally, such exposure can also result in spillback events at the human animal interface [66] and the spread of antibiotic resistance genes [67].

The “humanization” of microbiomes, that is, the acquisition of human-like microbiomes is recorded for captive mammals across taxa [37,38,40]. Bats inhabiting anthropogenic regions have a taxonomically and compositionally distinct microbiome as well [39]. We did not analyse samples across a habitat gradient and thus cannot look at how anthropogenic pressure affects bat microbiomes, but our study establishes urban baselines from South India. Future studies investigating the effects of anthropogenic pressure including pollution on microbiomes by sampling across a wilderness-rural-urban gradient are required, along with studies looking at their effects on hosts.

Although many of the roosts in our study were sampled repeatedly over several seasons (Table 1), these were opportunistic in nature and may be insufficient to investigate temporal dynamics of microbial abundance. However, we obtained similar taxonomic composition across samples from different time points (Figure 2a). This suggests temporal homogenisation, especially because the microbiome responds to consumed food [68]. This could be a consequence of our study region’s aseasonal climate with similar forage availability year-round. Moreover, our results except for alpha diversity metrics (Figure 4, Supplementary figure S6) display similar trends before and after pooling. While the former represents guano from a single roost collected on a given date, the latter is a time-averaged, roost-level sample. This suggests that our results could be extrapolated and generalised in a time-independent manner. Most existing studies have focussed on seasonal life-history events that change the microbiome in temperate species [52,69,70,71], and it is poorly understood how tropical, relatively aseasonal environments and periodic life-history events interact to affect species microbiomes.

## Conclusion

In conclusion, our study establishes baselines on microbiome composition and function from bat species common around human inhabitation in South India. We found microbiomes to be taxonomically but not functionally diet-dependent, and to poorly correlate with bat phylogeny. The presence of human pathogens and Cyanobacteria (also found in algal blooms in eutrophic water) in bat microbiomes suggests anthropogenic influences on them. We strongly recommend monitoring the microbiomes of urban wildlife in India in order to better understand its effects on hosts, and to monitor the spread of zoonoses and antibiotic resistance genes.

## Methods

### Species selection and panel creation

We used several methods to identify bat species from single-species roosts around Bengaluru city. *Pteropus medius* and *Cynopterus sphinx* were identified visually, and *Hipposideros speoris, H. lankadiva, and Lyroderma lyra* were identified using morphology and echolocation calls. To identify the single *Pipistrellus* roost which could not be identified to species level, we sequenced the cytochrome b (Cyt b) gene (which is commonly used for species-level identification in bats) and were able to identify it till the genus level.

We obtained 136 samples of six bat species from populations across an urban landscape from the city of Bengaluru, Karnataka, India (Figure 1). After pooling samples of the same roost collected on the same date, we finally obtained 39 different samples with a total of ten unique species-roost combinations collected over 20 months (Table 1).

### Sample collection and DNA extraction

We collected guano following a non-invasive ‘under the roost’ sampling protocol modified from [72]. All roosts sampled were single-species roosts, except for the *Hipposideros speoris* roost at site 6 which was transiently occupied by a few *Hipposideros lankadiva* individuals. At each roost site, we used ten 1 m × 1 m or 1 m × 2 m plastic sheets spread and pegged down under roost trees. We waited for two hours after placing down plastic sheets to wait for fresh guano to accumulate on them for frugivorous bats. For insectivorous bats, we collected samples (1 vial per quadrant and ∼200 mg of guano). We collected the faecal samples from each sheet using a sterile polyester swab. Samples were immediately placed in 1 mL DNA/RNA Shield (Zymo Research Corporation) to ensure sample inactivation at the field site, followed by agitation by swirling and scraping the sides to ensure that the samples were completely submerged by the buffer.

DNA was extracted using the Qiagen DNeasy Blood and Tissue Extraction Kit with the manufacturer-prescribed protocol used after minor modification. Briefly, guano samples were subjected to beat beating using Zirconia beads (0.5mm diameter, Biospec #11079105Z), ensuring their complete disruption to release nucleic acids. 20 µL of proteinase K was added to each sample, followed by vortexing and incubating in a heat block at 56° C overnight before lysis, binding, washing, and elution as listed in the manufacturer’s protocol. Field sample collection and laboratory protocols were approved by The National Centre for Biological Sciences Institutional Biosafety Committee (TFR:NCBS:36IBSC2021/UR1) and the Review Committee on Genetic Manipulation, Department of Biotechnology (BT/IBKP/035/2019).

### 16S rRNA library preparation and sequencing

Library preparation, sequencing and demultiplexing were carried out at the Next Generation Genomics facility at the National Centre for Biological Sciences – TIFR, Bengaluru. For library preparation, 40 µL of DNA extract was purified using 2.5x in-house carboxyl group-coated magnetic purification beads to a final volume of 5 µL. As described previously [73], N(1-10) spacer-linked primers were used to amplify the V3-V4 region of the bacterial 16S rRNA region. 10 µM of forward and reverse primers (2.5 µL each) were used in a 25 µL PCR reaction with Qiagen multiplex mastermix (12.5 µL, catalogue number 206145), and 5 µL of purified DNA. This reaction mix was initially denatured at 95° C for 15 minutes followed by 26 cycles of 94° C for 30 seconds, 55° C for 90 seconds, 72° C for 60 seconds, and with a final extension at 72° C for 10 minutes. PCR products were purified using 0.8x purification beads (prepared in-house) to a final volume of 22.5 µL. Amplicon sizes for random samples were checked using high sensitivity D1000 screen tape on a Tapestation 4200 (Agilent Technologies) for quality control. 3 µL (15-20 ng/µL) of purified PCR products were indexed in a 50 µL PCR reaction with Illumina nextera XT v2 indexes (Illumina) (5 µL of each index and 12.5 µL Qiagen hotstart mastermix). PCR conditions for this were the same as the PCR carried out for the 16S rRNA amplification, but for 8 cycles. Libraries were pooled and normalised to a final concentration of 2 nM and sequenced on an Illumina Miseq platform with a sequencing read length of 2x300.

### Sequence data processing and analyses

QIIME2 [74] was used for all sequence data-handling. Demultiplexed, trimmed, fastq files containing the raw reads were imported into QIIME2 in the Paired End Fastq Manifest Phred 33V2 format. Sequences were filtered by q-score using default settings (q-score value = 4). Next, we used the Deblur [75] algorithm set to a trim length of 190 bases for both forward and reverse reads based on quality scores, so chosen because quality began declining sharply after this value. Deblur groups sequences into Amplicon Sequence Variants (ASVs), which were used for all further analyses. Samples collected from the same roost and on the same date represent a colony-level averaged sample, and these were used for all analyses. Samples collected from the same roost on different dates could represent a different set of bats. Therefore, we performed all analyses using the samples directly, as well as after pooling ASVs from a given roost and collected on different dates in order to ensure independence. Pooling was done by directly summing up the ASV frequency values generated by Deblur. This results in a single time averaged, roost-level sample for each roost.

Rarefaction curves were plotted as a function of sequencing depth in QIIME2 after setting a maximum depth of 10,000 and plotted in Python ver. 3.12.2 (Supplementary figure S1). Except samples C2b, C2c, E1g and E1m, all sequences had adequate depth. One sample had a maximum depth < 10,000 ASVs and was not included in further analyses. We then used the greengenes2 [76] dataset to taxonomically classify our ASVs. Bar plots representing relative abundance of taxonomic units were plotted in R ver. 4.4.1 using the packages “tidyverse” [77] and “qiime2R” [78]. Principal Coordinates Analysis (PCoA) results were generated in QIIME2 using the “diversity” plugin and were plotted in R using the packages “tidyverse” and “qiime2R”.

To detect reported plant, animal, or zoonotic pathogens in our samples, we used the reference database used by the Multiple Bacterial Pathogen Detection (MBPD; accessed 28 August 2024) pipeline [79] and filtered it for pathogens found in our samples after greengenes2 classification. Greengenes2 classification lists the number of ASVs classified per taxon, and the MBPD database uses the Greengenes classification scheme for analyses as well. We therefore subset our samples for pathogenic taxa reported in the MBPD database.

A neighbour-joining tree was generated in R using the Weighted UniFrac distance matrix from pooled samples using the packages “ape [80] and “phangorn” [81,82].

### Diversity metric calculations

We generated per-sample metrics of diversity (alpha diversity) as well as cross-sample metrics of diversity (beta diversity) in QIIME2 using the “diversity” plugin. For alpha diversity metrics, we calculated the Shannon diversity index, Faith Phylogenetic Diversity (PD) [83], and Pielou’s evenness [84] for each sample. For beta diversity metrics, we calculated Jaccard, Bray–Curtis, and Weighted Unifrac metrics across pairs of samples. Results from these were plotted in Python using the “seaborn” [85] library. Two sided t-tests for comparing alpha and beta diversity metrics with diet type were performed using the Python library “scipy” [86].

### Functional annotation and analyses

We used PICRUSt2 [87] for predicting functional abundances. PICRUSt2 outputs predicted abundances of enzymes classified under the Enzyme Commission (EC), and alternatively, KEGG Orthology (KO) naming schemes. To visualise this, the abundances of enzymes classified as per the EC numbering scheme were used from the file titled “pred_metagenome_unstrat.tsv” output by PICRUSt2. These were first converted into biom format and then imported into QIIME2 in the “FeatureTable[Frequency]” format. After this, we computed the Jaccard dissimilarity metric between samples using the “diversity-lib” plugin and performed PCoA analyses using the “diversity” plugin in QIIME2. Results from this were visualised in R using the packages “tidyverse” and “qiime2R”. Principal Components Analysis (PCoA) plots from the KO predicted pathway abundances output by PICRUSt2 were visualised in R ver. 4.3.1 using the package “ggpicrust2”.

## Declarations

### Ethics approval and consent to participate

Ethics approvals obtained were IBSC: TFR:NCBS:36IBSC2021/UR1 and from the Review Committee on Genetic Manipulation, Department of Biotechnology (BT/IBKP/035/2019). IBSC and RCGM approvals were obtained in February 2021 and December 2021, respectively and they are valid till the end of the project.

### Consent for publication

Not applicable.

### Data availability

All intermediate files and codes are available at the Zenodo url https://doi.org/10.5281/zenodo.15909932

Sequence data are available at the BioProject accession PRJNA1271418.

### Authors’ contributions

BC and UR conceptualization; DS, AS, and ABR generated data, VI performed the analyses with advice from BC; VI and BC prepared the original draft with inputs from ABR; VI prepared the figures; All authors reviewed the manuscript.

## Acknowledgements

We are grateful to Rajesh Puttaswamaiah from Bat Conservation India Trust for his invaluable support in locating insectivorous bat roosts in and around Bangalore. We thank Tikily Tayeng, Fathima Aslaha and Shraddha Kumari for help with the sampling. We also acknowledge the support of the NCBS next-generation genomics facility and biosafety facilities. BC thanks Centre for Climate Change and Sustainability and Trivedi School of Biosciences, Ashoka University for funding support. BC thanks Kritika M. Garg for her help in data analyses and for commenting on an earlier version of the manuscript.

