## Supplementary material for "Anthropogenic habitats shape gut microbiome composition in Southern Indian bats": Supplemetary information

Supplementary information

### S1: Sampling coverage

Rarefaction curves for each sample (unpooled) displaying the number of ASVs at each sampling depth, plotted per species. The maximum depth is set to 10,000.

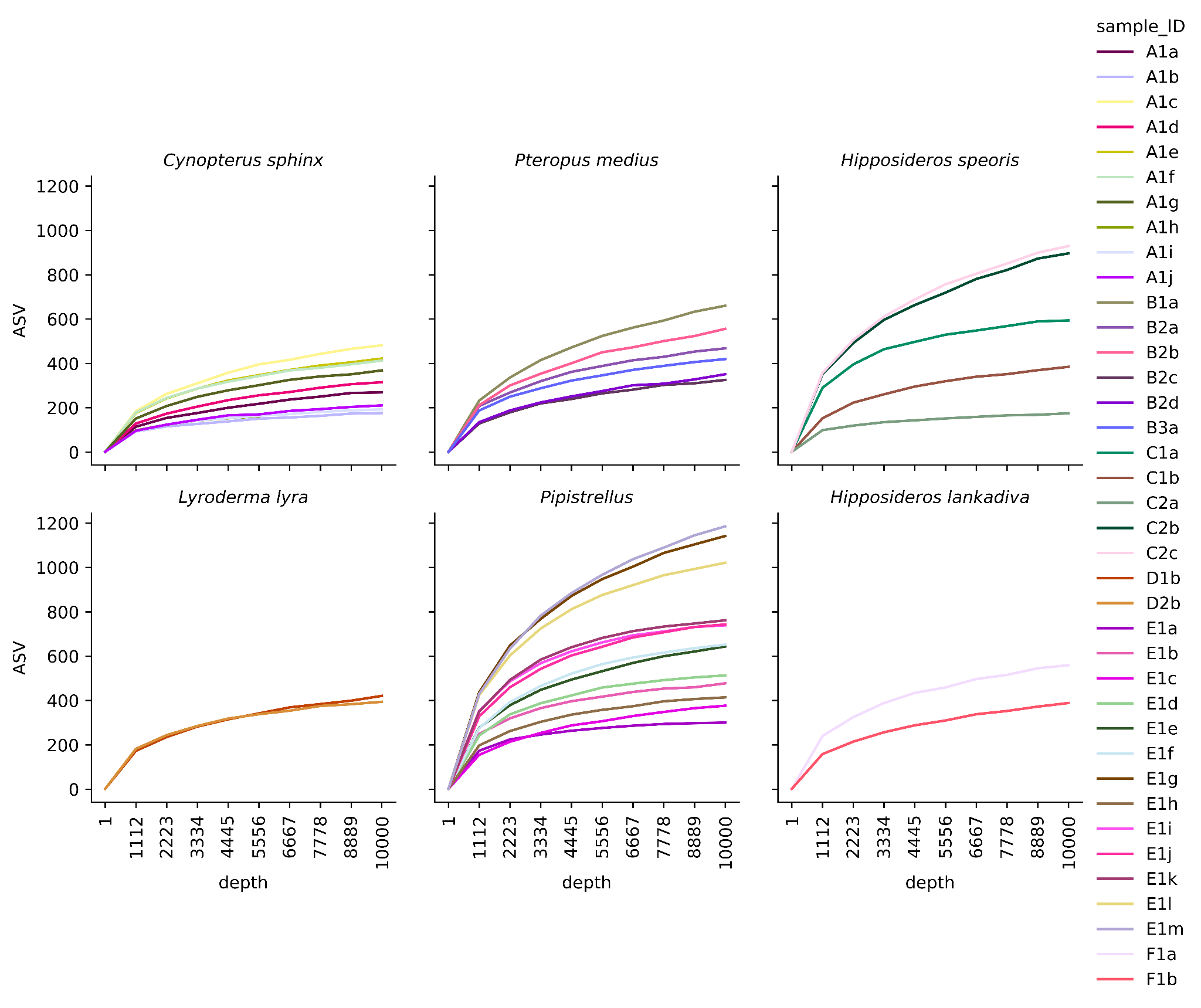

### S2: Stacked taxonomic bar plots

Stacked bar plots displaying the proportion of various taxa in each sample at the (a) kingdom, (b) phylum, and (c) class levels. The 10 most abundant taxa at each level are displayed by name in (b) and (c). The x-axes represent sample codes.

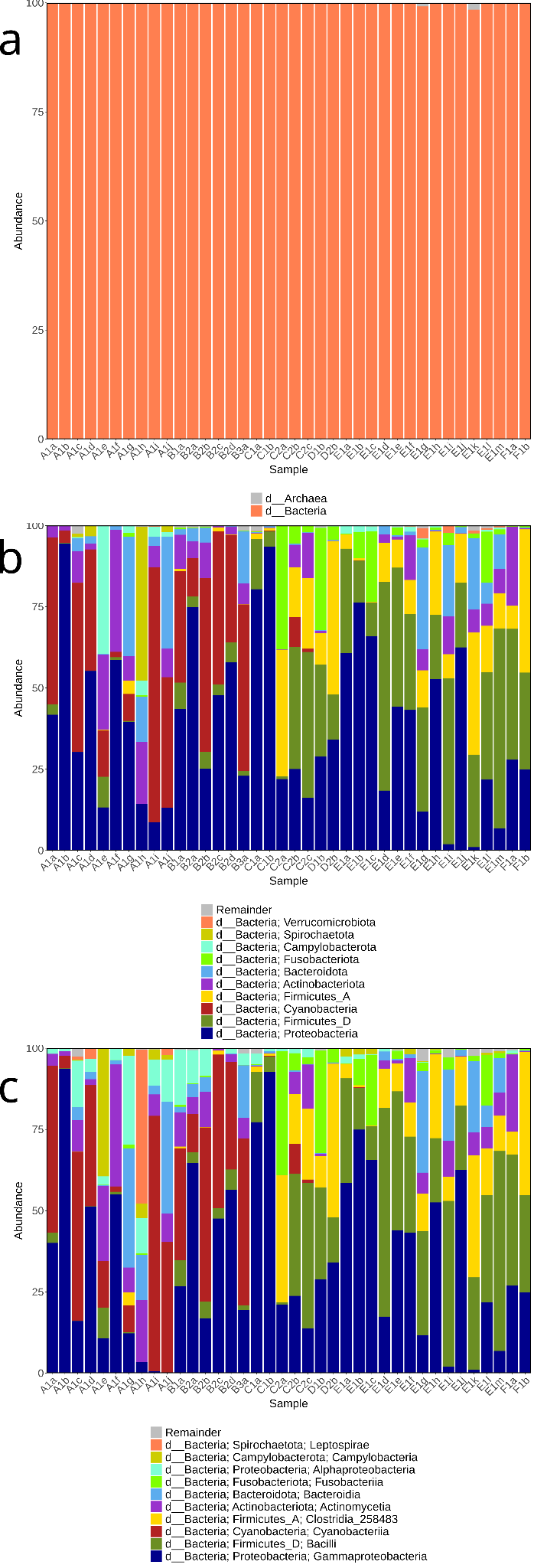

### S3: Sample compositional relatedness amongst unpooled samples:

### Principal Coordinates Analysis plots of unpooled samples using various distance metrics and microbiome taxonomy and coloured by species identity. Each point represents a sample collected on a given date from a given roost. Points with the same colour and shape represent samples collected from the same roost on different dates.

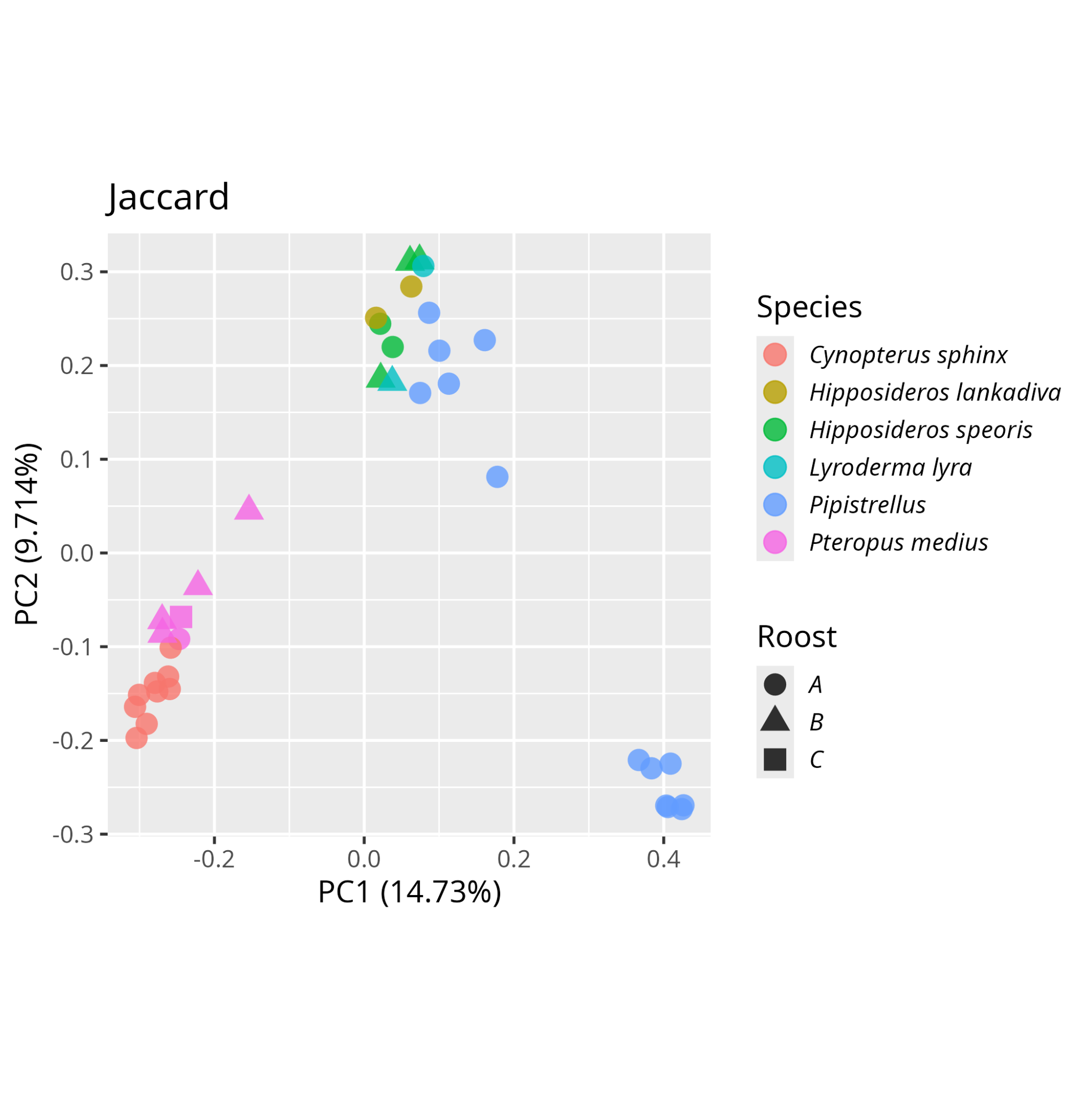

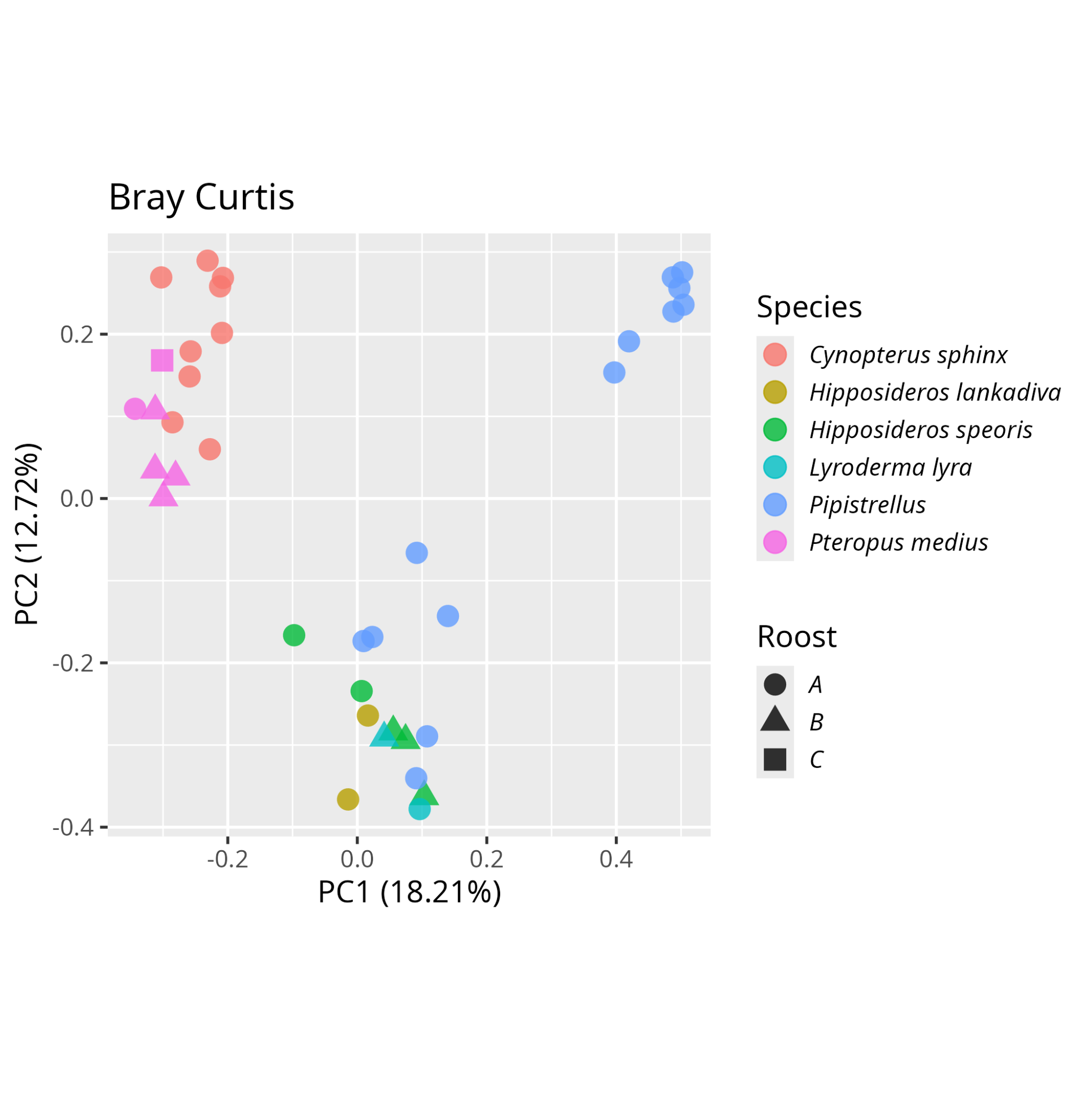

### S4: Sample compositional relatedness amongst pooled samples:

### Principal Coordinates Analysis plots of pooled samples using various distance metrics and microbiome taxonomy and coloured by species identity. Each point represents a sample collected on a given date from a given roost. Points with the same colour and shape represent samples collected from the same roost on different dates.

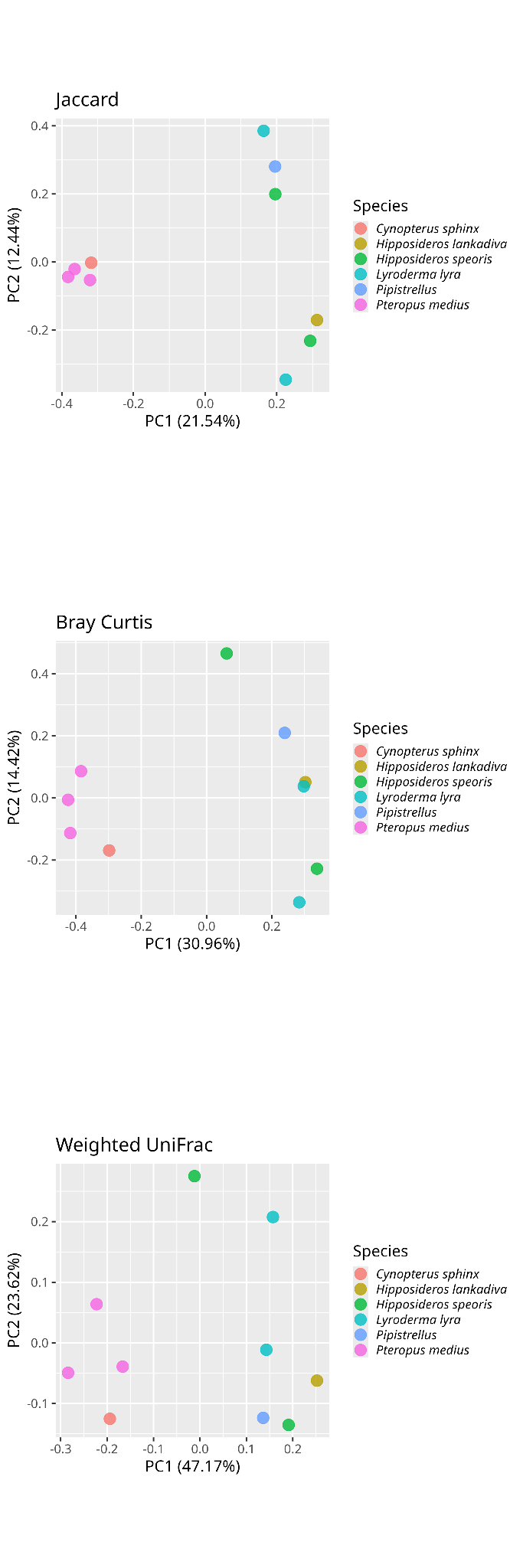

### S5: Functional relatedness amongst pooled samples

Above: Principal Coordinates Analysis plot of pooled samples using Jaccard distance metric and predicted enzyme abundances as per the Enzyme Commission (EC) and coloured by species identity. Each point represents a sample collected on a given date from a given roost. Points with the same colour and shape represent samples collected from the same roost on different dates. Below: Principal Components Analysis plot of pooled samples of predicted pathway abundances using KO pathway numbers. Ellipses indicate 95% confidence ellipses. Density plots are plotted on the axes.

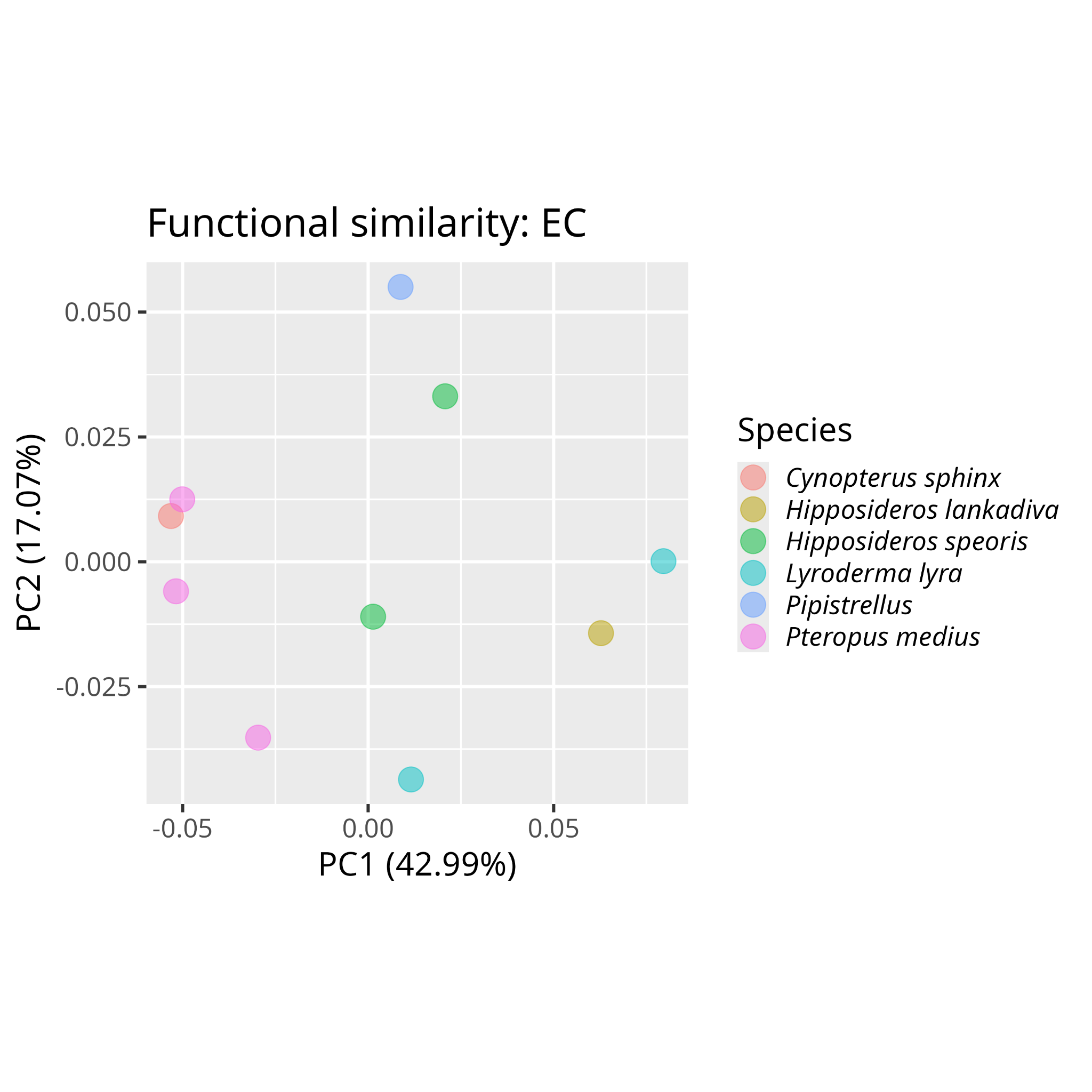

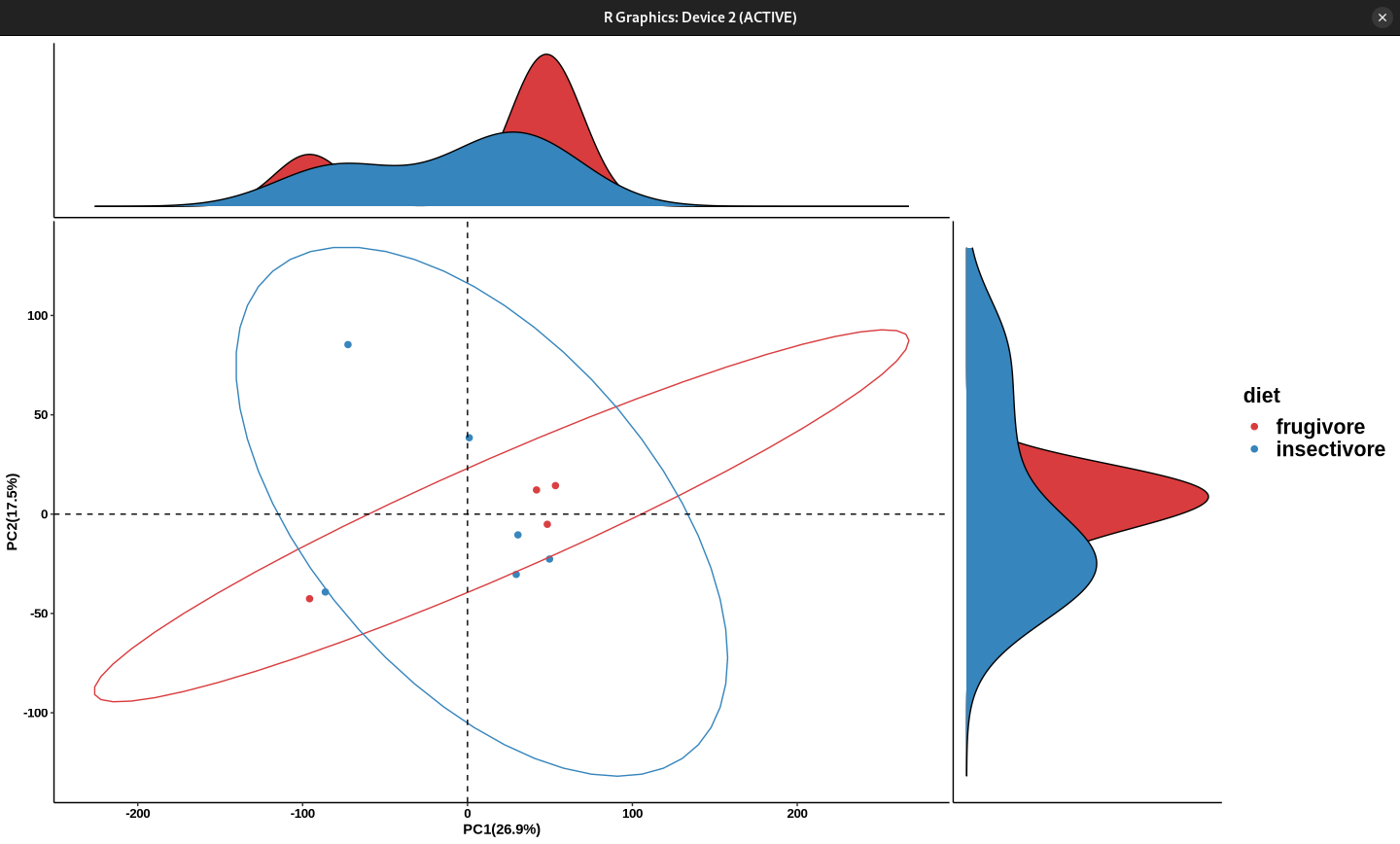

### S6: Alpha diversity metrics amongst pooled samples

Strip plots of Pielou Evenness, Faith’s PD, and Shannon Entropy for each diet type. t and p values represent values from two-sided t-tests.

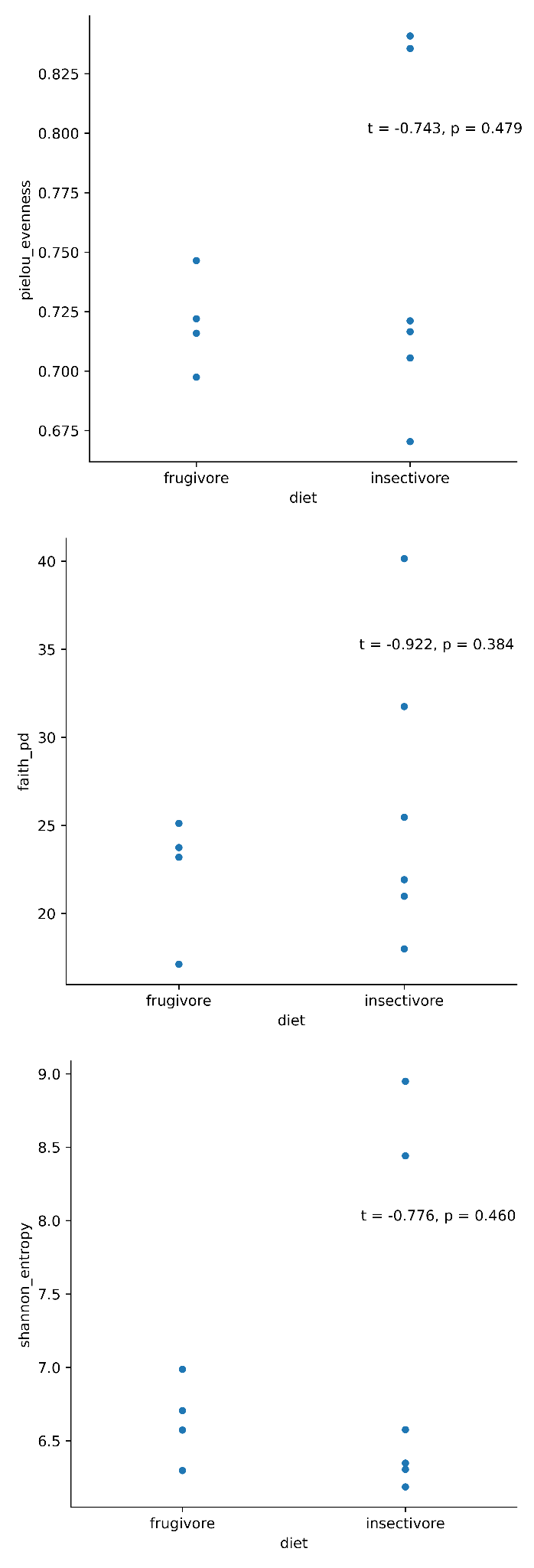

### S7: Distances to samples: unpooled data

A boxplot using unpooled data of the distances of each datapoint to other data points of the same or different diet type. All data points are overlaid in blue. Top: Jaccard distance metric. Bottom: Bray – Curtis distance metric.

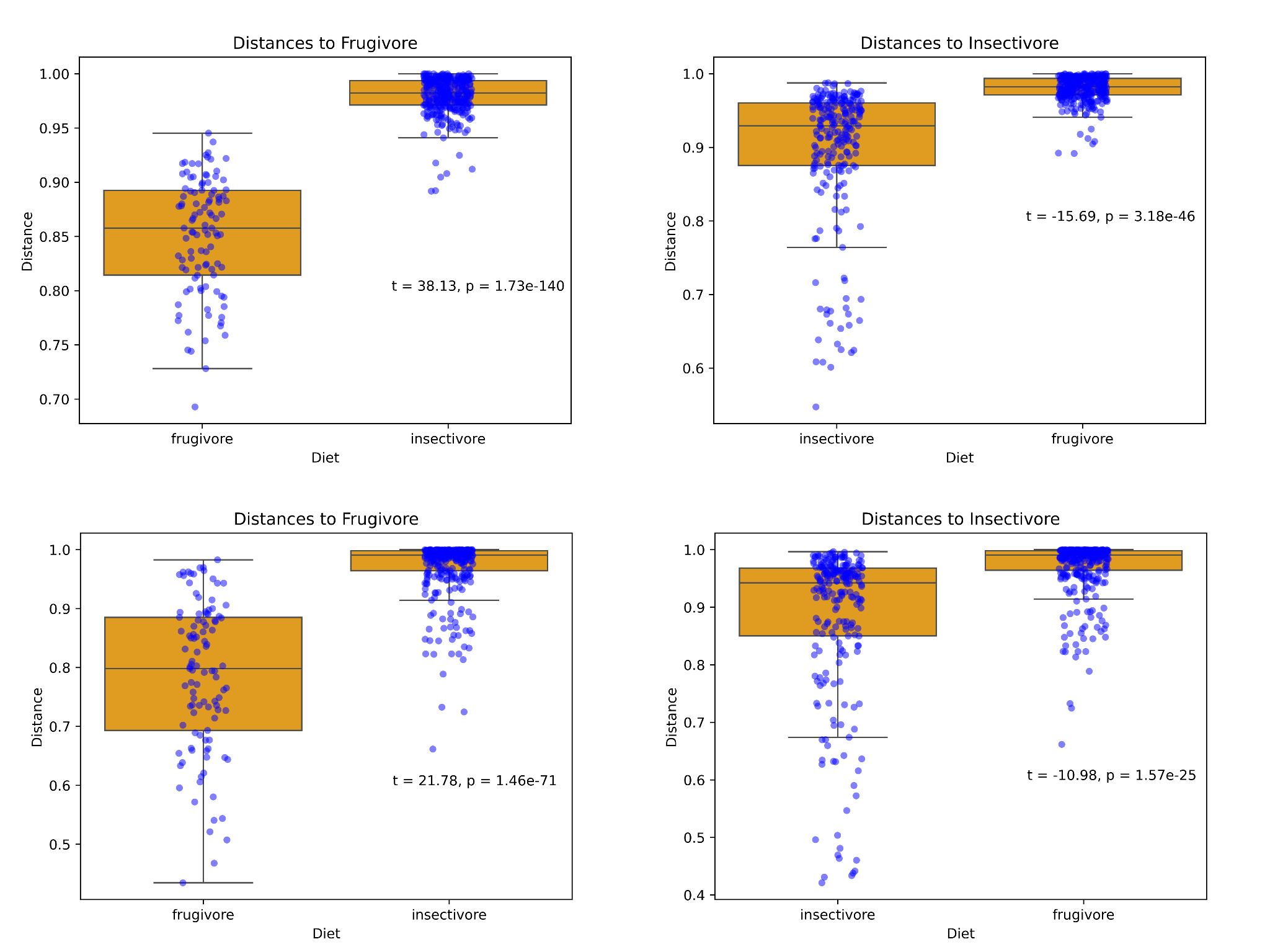

### S8: Distances to samples: pooled data

A boxplot using pooled data of the distances of each datapoint to other data points of the same or different diet type, using various distance metrics. All data points are overlaid in blue. Top: Jaccard distance metric. Middle: Bray – Curtis distance metric. Bottom: Weighted UniFrac distance metric.

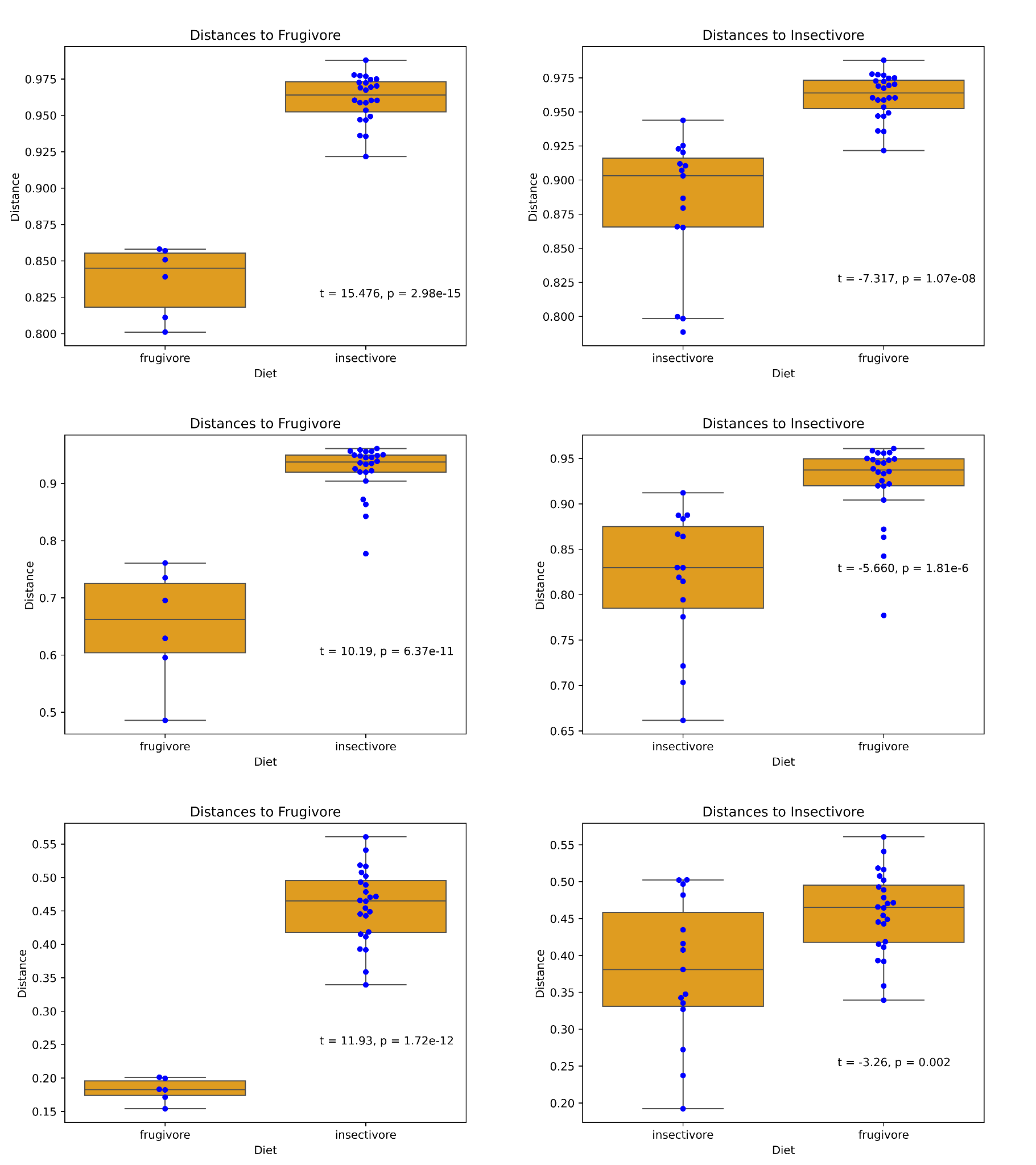
